# Non-additive effects of changing the cytochrome P450 ensemble: incorporation of CYP2E1 into human liver microsomes and its impact on CYP1A2

**DOI:** 10.1101/685545

**Authors:** Nadezhda Y. Davydova, Bikash Dangi, Marc A. Maldonado, Nikita E. Vavilov, Victor G. Zgoda, Dmitri R. Davydov

## Abstract

The aim of this study is to investigate the ability of ethanol-inducible CYP2E1 to interact with other cytochrome P450 species and affect the metabolism of their substrates. As a model system we used CYP2E1-enriched microsomes obtained by incorporation of purified CYP2E1 protein into HLM. Using the method based on homo-FRET in homo-oligomers of CYP2E1 labeled with BODIPY 577/618 maleimide we demonstrated that the interactions of CYP2E1 with microsomes result in dissociation of the protein homo-oligomers. The finding that this effect is much better pronounced in HLM as compared to the microsomes containing no P450 proteins indicates the formation of mixed oligomers of CYP2E1 with other P450 species that takes place in expense of dissociation of the homo-oligomers.

Incorporation of CYP2E1 into HLM results in a multifold increase in the rate of metabolism of CYP2E1-specific substrates *p*-Nitrophenol (pNP) and Chlorzaxozone (CLZ). The rate of their oxidation remains proportional to the amount of incorporated CYP2E1 up to the content of 0.3-0.4 nmol/mg protein (or about 50% CYP2E1 in the P450 pool). These results demonstrate that the incorporated CYP2E1 becomes a fully-functional member of the P450 ensemble and do not exhibit any detectable functional differences with the endogenous CYP2E1 in HLM.

Enrichment of HLM with CYP2E1 results in a pronounced alteration of the metabolism of 7-etoxy-4-cyanocoumarin (CEC), the substrate of CYP2C19 and CYP1A2, that suggests an important augmentation of the involvement of CYP1A2 in its metabolism. This effect goes together with a remarkable increase in the rate of dealkylation of CYP1A2-specific substrate 7-ethoxyresorufin by CYP2E1-enriched HLM. Furthermore, probing the interactions of CYP2E1 with model microsomes (Supersomes™) containing individual P450 enzymes we found that CYP2E1 efficiently interacts with CYP1A2, but lacks any ability to form complexes with CYP2C19. This finding goes inline with CYP2E1-induced redirection of the main route of CEC metabolism from CYP2C19 to CYP1A2.

---

The functional properties of the drug-metabolizing system of human liver are largely determined by the properties of the ensemble of cytochromes P450, which is responsible for metabolism of over 75% of all marketed drugs and new drug candidates [1-3]. It is commonly recognized that the inter-individual, age-dependent and temporal variations in drug metabolism are largely dictated by the changes in the composition of the ensemble of multiple drug-metabolizing cytochrome P450 species. However. recent studies on a large selection of human liver microsomes (HLM) demonstrated the lack of a straight correlation between the composition of human cytochrome P450 ensemble and the profile of drug metabolism [4]. At the same time, there is an emerging recognition of the functional importance of physical interactions between multiple P450 species co-localized in the microsomal membrane [5-9]. Due to complex effects of these interactions any change in the content of a particular P450 enzyme may affect drug metabolism in a complex, hard-to-predict manner. Therefore, a systematic exploration of the network of P450-P450 interactions and identification of their functional consequences represents an issue of emerging importance.

Of particular practical significance are the functional effects of the changes in the composition of P450 ensemble induced by alcohol consumption. Although the multi-fold increase in the content of cytochrome P450 2E1 (CYP2E1) in liver observed in both alcoholics and moderate alcohol consumers is well documented [10-14], the role of CYP2E1 in the instances of alcohol-drug interactions is commonly considered insignificant due to a minor role of this enzyme in drug metabolism. However, the involvement of CYP2E1 in alcohol-drug interactions may stretch far beyond its immediate effects on the metabolism of its substrates.

There is a limited amount of information on possible functional effects of CYP2E1 on other human P450 species. In an early study with model microsomes Tan and co-authors demonstrated mutual inhibitory effects of CYP2E1 and CYP2A6, which were attributed to a competition for NADPH-cytochrome P450 reductase [15]. In the same study the authors also observed a potentially important effect of co-expression of CYP2E1 with CYP2A6 on H_2_O_2_ production by CYP2A6. While NADPH-dependent generation of H_2_O_2_ by CYP2A6 was drastically increased in the presence of coumarin, addition of this CYP2A6 substrate results in decreased H_2_O_2_ production by the system composed of CYP2A6, CYP2E1 and CPR [15]. In a latter study in a micellar reconstituted system Kelley and co-authors explored the reciprocal effects of rabbit CYP1A2 and CYP2E1 [16]. The authors demonstrate that, while the addition of CYP1A2 does not affect CYP2E1-dependent metabolism of N-nitrosodimethylamine, addition of CYP2E1 considerably intensifies CYP1A2-dependent dealkylation of 7-ethocyresorufin (ERR). Based on the observation that this effect was most pronounced at sub-saturating concentrations of CPR the authors concluded that the stimulating effect of CYP2E1 on CYP1A2 may be exerted through the formation of the CYP1A2-CYP2E1 complexes where the interactions of CYP1A2 with CPR are promoted [16].

In our study of interactions between CYP2E1 and CYP3A enzymes in model microsomes we demonstrated the formation of the complexes of CYP2A1 with either CYP3A4 or CYP3A5, where the activity of CYP2E1 is boosted, while the functionality of the CYP3A enzymes remains unaffected [17]. In our recent study we also explored the interactions between CYP2E1 with CYP2D6 and observed the formation of their heteromeric complexes in microsomal membrane. We demonstrated that these interactions inhibit the activity of CYP2E1, while exerting a time-dependent activating effect on CYP2D6 in the presence of its specific substrates, 3-[2-(N,N-diethyl-N-methyl ammonium)ethyl]-7-methoxy-4-methylcoumarin (AMMC) and bufuralol [9]. Of particular importance might be the effect of CYP2E1 on the stoichiometry of futile cycling and substrate oxidation by CYP2D6 revealed in a decrease in the electron leakage from CYP2D6 through the peroxide-generating pathways [9]. This effect is reminiscent of an impact of CYP2E1 on H_2_O_2_ production by CYP2A6 observed by Tan and co-workers [15].

All these results demonstrate complex functional relationship between CYP2E1 and other drug-metabolizing P450 species which deserves detailed attention in view of its potential involvement in alcohol-drug interactions. Of particular interest is possible crosstalk between CYP2E1 and CYP1A2 evidenced in the study of Kelley and co-authors [16]. The effect of ethanol-induced increase in CYP2E1 concentration in liver on CYP1A2 may be involved in pronounced interactions of alcohol with such drug substrates of CYP1A2 as tricyclic antidepressnts (amitryptiline, clomipramine, imipramine, etc.) [18-19]. Most of the drugs of this type are also metabolized by CYP2C19 and possible rerouting of drug metabolism between the two enzymes may also play a role in their interactions with alcohol.

The aim of the present study is to probe the interactions of CYP2E1 with cytochromes P450 present in HLM and explore the effect of these interactions on the functional properties of CYP1A2 and CYP2C19. In a search for a model system suitable for these studies we recently introduced a method based on incorporation of purified cytochromes P450 into microsomal membrane [9, 17, 20]. An important characteristic of this strategy is its adaptability for incorporating genetically or chemically modified proteins. This includes the incorporation of proteins bearing fluorescent probes and light-activating crosslinkers. Therefore, this strategy may be used for monitoring protein-protein interactions in HLM with various biophysical techniques, as well as for identifying the P450 interactions partners with the technique of molecular fishing. Results of the present study demonstrate high utility of this approach in the studies of interactions between multiple cytochrome P450 species and their effect on drug metabolism. Using this strategy we evidenced the formation of the complexes of CYP2E1 with other P450 species and demonstrated the effects of CYP2E1 on the activity of CYP1A2, the enzyme involved in metabolism of a wide variety of commonly used drugs known for their pharmacokinetic interactions with alcohol.

## Materials and Methods

***Materials.*** *p*-Nitrophenol, *p*-Nitroanizol, and DL-dithiothreitol (DTT) were the products of Sigma-Aldrich (St. Louis, MO). NADPH tetrasodium salt was from EMD Millipore (Billerica, MA). 7-ethoxyresorufin (ERR) was from AnaSpec (San Jose, CA, USA). 7-etoxy-4-cyanocoumarin (CEC), 7-hydroxy-4-cyanocoumarin (CHC) and BODIPY 577/618 maleimide were from Invitrogen/Molecular Probes Inc. (Eugene, CA).. Chlorzoxazone, 6-hydroxy chlorzoxazone and resorufin were the products of Cayman Chemical (Ann Arbor, MI). All other reagents were of ACS grade and used without additional purification.

***Protein expression and purification.*** N-terminally truncated (Δ3-20) and C-terminally His-tagged CYP2E1 [21] was expressed in *E. coli* and purified following the procedure similar to that described for CYP2B6 [22] followed by ion-exchange chromatography on Macro-Prep CM Support resin (Bio-Rad Laboratories, Hercules, CA).

***Microsomes from insect cells bearing recombinant human cytochromes P450 (SupersomesTM)*** were the products of BD Gentest, now a part of Corning Life Sciences (Tewksbury, MA). In the present study we used the preparations containing CYP2E1 (SS2E1, lot 44748), CYP1A2 (SS1A2, lot 69375), CYP2B6 (SS2B6, lot 31487), CYP2C8 (SS2C8, lot 8239002), CYP2C9 (SS2C9, lot 41274), CYP2C19 (SS2C19, lot 73445), CYP2D6 (SS2D6, lot 7114001) and CYP3A4 (SS3A4, lot 35933). All those preparation contained human NADPH-cytochrome P450 reductase (CPR) and cytochrome b5, except for SS2C8 and SS1A2, which had no cytochrome *b*_5_ co-expressed. In the experiments on CYP2E1 interactions with microsomal membrane we also utilized Supersomes™ containing rat CPR (SS-RR, lot 5274006) or human CPR and cytochrome b5 (SS-B5, lot 41774) as control membranes bearing no cytochromes P450.

***Preparations of human liver microsomes (HLM).*** In this study we used two different preparations of pulled human liver microsomes. The preparation refered here as HLM-1 was prepared by differential centrifugation from the sample of human liver S9 pool (lot number 3212595) obtained from BD Gentest. The HLM sample refered here as HLM-2 is an InVitroCYP™ M-class 50-donor mixed gender pooled HLM preparation (lot LFJ) obtained from BioIVT corporation (Baltimore, MD).

***Incorporation of CYP2E1 into HLM*** was performed by incubation of undiluted suspensions of HLM (20-25 mg/ml protein, 10-13 mM phospholipid) in 125 mM K-Phosphate buffer containing 0.25M Sucrose with purified CYP2E1 for 16 - 20 hours at 4°C at continuous stirring. CYP2E1 was added in the amount ranging from 0.25 to 2 molar equivalents to the endogenous cytochrome P450 present in HLM. Following the incubation the suspension was diluted 4-8 times with 125 mM K-Phosphate buffer, pH 7.4 containing 0.25 M sucrose and centrifuged at 53,000 rpm (150,000 g) in an Optima TLX ultracentrifuge (Beckman Coulter Inc., Brea, CA, USA) with a TLA100.3 rotor for 90 min at 4 °C. The pellet was resuspended in the same buffer to the protein concentration of 15-20 mg/ml. The amount of incorporated cytochrome P450 was calculated from the difference between the heme protein added to the incubation media and the enzyme found in the supernatant. According to the results of this assay our procedure resulted in incorporation of 96-98% of the added protein into the microsomal membrane.

***Determinations of protein and phospholipid concentrations in microsomal suspensions*** were performed with the bicinchoninic acid procedure [23] and through the determination of total phosphorus in a chloroform/methanol extract according to Bartlett [24], respectively.

***The concentration of NADPH-cytochrome P450 reductase*** in microsomal and proteoliposomal membranes was determined based on the rate of NADPH-dependent reduction of cytochrome *c* at 25ºC [25] monitored by an increase in the difference in absorbencies at 550 and 541 nm and calculated using the differential (“reduced-minus-oxidized”) extinction coefficient of cytocrome *c* of 18 mM^−1^cm^−1^ [26]. Effective molar concentration of CPR was estimated using the turnover number of 3750 min^−1^, which we determined with purified CPR quantified from its absorbance at 456 nm using the extinction coefficient of 21.4 mM^−1^ cm^−1^ [27].

***The total concentration of cytochrome P450 in HLM*** was determined with a variant of “oxidized CO versus reduced CO difference spectrum” method [28]. In our assay we used Microsome Solubilization Buffer (MSB) composed of 100 mM potassium phosphate pH 7.3, 10% Glycerol, 0.5% Sodium Cholate and 0.4% Igepal CO-630. Prior to addition of the microsomal suspension the buffer was saturated with CO through bubbling the gas for 40-60 sec. 100 µl of CO-saturated MSB were placed into a quartz microcuvette. After recording a baseline in 380-600 nm region the cell was supplemented with 5µl of HLM sample (10-20 mg/ml) and the spectrum was recorded. Reduction was achieved by addition of a small amount of dithionite dust. The process of reduction was followed through monitoring the changes in the difference of optical densities (DOD) at 450 and 490 nm. The spectrum of absorbance of the reduced sample was taken when the DOD stabilizes. The difference between the spectra taken after and before reduction was calculated and used to determine the concentration of cytochromes P450 using the extinction coefficient of 106 mM^−1^ cm^−1^ for DOD between the maximum of the peak around 450 nm (448-451 nm) and the plateau at 490 nm.

***Determination of the content of cytochrome b5 in HLM*** was based on its NADH-dependent reduction and performed following the procedure described by Kennedy [29].

***Mass-spectrometric analysis of the content of cytochromes P450 and NADPH-cytochrome P450 reductase in HLM*** was performed with a triple quadrupole mass spectrometer using the method of multiple reaction monitoring (MRM). For each protein, one standard peptide with three transitions was used. The peptides selected were arranged into one SRM assay. The information (m/z of precursors, m/z of transition ions, CE values, b, y ions transition ions and MS platform were used for the analysis) on SIS and the target peptides is provided in Table S1 found in Supplementary Material. Separation of peptides from digested HLM samples was carried out using the UPLC Agilent 1290 system including a pump and an autosampler. The sample was loaded into the analytical column Eclipse Plus SBC-18, 2.1 × 100 mm, 1.8 um, 100 A. Peptide elution was performed by applying a mixture of solvents A and B. Solvent A was HPLC grade water with 0.1% (v/v) formic acid, and solvent B was 80%(v/v) HPLC grade acetonitrile/water with 0.1% (v/v) formic acid. The separations were performed by applying a linear gradient from 3% to 32% solvent B over 50 min, then from 32% to 53% solvent B over 3 min at 300μl/min followed by a washing step (5 min at 90% solvent B) and an equilibration step (5 min at 3% solvent B). Ten microliters of each sample were applied on chromatographic column. The quantitative analysis was performed using Agilent 6495 Triple Quadrupole (Agilent, USA) equipped with a Jet Stream ionization source. The following parameters were used for the Agilent Jet Stream ionization source: the temperature of the drying gas of 280°C, 18 psi pressure in the nebulizer, 14 L/min flow rate of the drying gas, and 3000V voltage on the capillary.

The standard samples of target peptides were obtained using the solid-phase peptide synthesis on the Overture (Protein Technologies, USA) according to the published method [30]. The isotope-labeled lysine (13C6,15N2) or arginine (13C6,15N4) was used for isotope-labeled peptide synthesis instead of the unlabeled lysine or arginine, correspondingly. Concentration of the synthesized peptides was quantified through acidic hydrolysis followed by analysis of derived amino acids with fluorimetric detection [31].

***Activity measurements with fluorogenic substrates*** The rate of O-dealkylation of CEC and ERR were measured with a real-time continuous fluorometric assay. A suspension of microsomes was added to 250 µl of 0.1 M Na-HEPES buffer, pH 7.4, containing 60 mM KCl to a final CPR concentration of 0.003 – 0.01 µM. The mixture was placed into a 5 × 5 mm quartz cell with continuous stirring and thermostated at 30 °C. An aliquot of a 15-20 mM stock solution of CEC or 0.4-2 mM stock solution of ERR in acetone was added to attain the desired concentration in the range of 0.5 - 250 µM (CEC) or 0.03 - 10 µM (ERR). The reaction was initiated by addition of 20mM solution of NADPH to the concentration of 200 µM. Increase in the concentration of products of dealkylation (CHC or resorufin) was monitored with a Cary Eclipse fluorometer (Agilent Technologies, Santa Clara, CA, USA) or a custom-modified PTI QM-1 fluorometer (Photon Technology International, New Brunswick, NJ) [9]. With both instruments the emission wavelength was set to 455 and 585 nm for the measurements with CEC and ERR, respectively. The excitation was performed with a monochromatic light centered at 405 or 560 nm (for CEC and ERR, respectively) with 5 nm bandwidth (Cary Eclipse) or with a light emitted by a 405 nm (CEC) or 532 nm (ERR) laser diode module (Thorlabs Inc, Newton, NJ. The rate of formation of the fluorescent products was estimated by determining the slope of the linear part of the kinetic curve recorded over a period of 3 – 5 min. Calibration of the assay was performed by measuring the intensity of fluorescence in a series of 5-10 samples of the same reaction mixture containing increasing concentrations of CHC or resorufin.

***Measurements of the rate of hydroxylation of p-nitrophenol*** was preformed with absorbance spectroscopy [32]. To measure the rate of formation of *p*-nitrocatechol (pNC) by HLM we mixed 3-5µl of the microsomal suspension with reaction buffer (0.1 M Na-Hepes buffer containing 60mM KCl) and a desired amount of 10-20 mM stock solution of pNP in reaction buffer to the total volume of 80 µl. After a preincubation for 1 min in a water bath at 30 °C the reaction was initiated by addition of 2µl of 20 mM NADPH solution. The probes were incubated at 30 °C for 30 min in a shaking water bath and the reaction was stopped by addition of 24 µl of 20% Trichloroacetic acid. The precipitated protein was removed by centrifugation at 15,000g for 2 min and the supernatant was supplemented with 16 µl of 5M NaOH. The spectra of absorbance of the samples in 450 – 700 nm region were recorded in a 100µl microcuvette (optical path 1 cm). As a baseline we used a spectrum of the sample treated as above and containing all ingredients except for pNP. To quantify the amount of the formed pNC and obtain the dependencies of the reaction rate on substrate concentration we subjected the series of spectra obtained with pNP concentrations increasing from 0 to 250 µM to the principal component analysis (PCA) procedure. The spectra of the first two principal components were approximated with a combination of the standard spectra of absorbance of pNP and pNC supplemented with a first or second order polynomial, which was used to compensate for possible variation in turbidity of the samples. The maximum of absorbance of pNC was found at 514 nm and the value of the extinction coefficient of 10.3 mM^−1^ cm^−1^ at 546 nm [32] was used to normalize the spectral standard to 1µM concentration. The concentration factors found in this fitting were used to scale the first two vectors of loading factors and combine them into a dependence of the concentration of formed pNC on the concentration of pNP.

***Activity measurements with chlorzoxazone*** were performed in 0.1 M Na-HEPES buffer, pH 7.4, containing 60mM KCl. The reaction was started by addition of NADPH to the final concentration of 0.5 mM and carried on for 10 min at 30 °C. Chlorzoxazone was added as a 25 mM solution in 60 mM KOH. Reaction was quenched with an addition of 25% of the incubation volume of 1 M solution of formic acid in acetonitrile containing a known concentration 3,5-dibromo-4-hydroxybenzoic acid as internal standard, followed by centrifugation at 9,300 g for 10 min.

The amount of formed 6-hydroxychlorzoxazone was quantified by liquid chromatography-mass spectrometry. An LC-20AD series high-performance liquid chromatography system (Shimadzu, Columbia, MD) fitted with a HTC PAL autosampler (LEAP Technologies, Carrboro, NC) was used to perform chromatography on a Kinetex® reverse-phase column (50 × 2.1 mm, 2.6 µm; Phenomenex, Torrance, CA). Chromatographic separation was performed using an isocratic method at the flow rate of 0.18 ml/min (mobile phase A) and 0.22 ml/min (mobile phase B) for 5 min. The quantification of the metabolite was conducted using an API 4000 Q-Trap tandem mass spectrometry system manufactured by Applied Biosystems/MDS Sciex (Foster City, CA) using turbospray ESI operating in negative ion mode. The mass spectrometer parameters were set at curtain gas, 20; collision gas, medium; ion spray voltage, −4500; ion source gas 1, 70; ion source gas 2, 50; temperature, 500, declustering potential, −550; entrance potential, −5; collision energy, −25, collision cell exist potential, 10. The analyte (6-hydroxychlorzoxazone) and the internal standard (3,5-dibromo-4-hydroxybenzoic acid) were detected using multiple reaction monitoring (MRM) mode by monitoring the *m/z* transition from 184.0 to 120 and 294.8 to 251.0, respectively. Determination of the product amounts was achieved with a calibration curve ranging from 0 to 2000 nM concentration of 6-hydroxychlorzoxazone.

***Normalization of the rates of substrate metabolism determined in HLM samples*** was performed based on the concentration of CPR determined form the rate of NADPH-dependent reduction of cytochrome *c* as described above. The rates of substrate metabolism by Supersomes™ were normalized on the concentration of cytochrome P450 (due to excess of CPR over cytochrome P450 in these preparations).

***Monitoring the interactions of CYP2E1-BODIPY with microsomal membranes*** was performed with the use of a Cary Eclipse spectrofluorometer equipped with a Peltier 4-position cell holder. The excitation of donor phosphorescence was performed with monochromatic light centered at 505 nm with 20 nm bandwidth. Alternatively the measurements were done with the use a custom-modified PTI QM-1 fluorometer (Photon Technology International, New Brunswick, NJ) equipped with a thermostated cell holder and a refrigerated circulating bath. In this case the excitation at 505 nm was performed with an M505L3 LED module (Thorlabs Inc, Newton, NJ). The spectra in the 570 – 800 nm wavelength region were recorded repetitively with the time interval varying from 0.5 to 15 min during the course of monitoring (5 – 16 hours). The experiments were performed at continuous stirring at 4 °C in 100 mM Na-Hepes buffer containing 150 mM KCl and 250 mM sucrose.

***Calculation of the surface density of cytochromes P450 in microsomal membrane*** was based on the molar ratio of microsomal phospholipids to cytochromes P450 (*R*_L/P_). In these calculations we used the value of 0.95 nm^2^ for the area of microsomal membrane corresponding to one molecule of phospholipid in a monolayer [33], similar to the approach used in our earlier reports [9, 17, 20]

***Data analysis.*** Analysis of series of spectra obtained in fluorescence spectroscopy experiments was done by principal component analysis (PCA) [34], which was used as a method of global analysis that minimizes the contribution of incidental variation in the shape of the spectra and makes the results more robust to a minor displacement of the emission band (a blue shift of approx. 2 nm) that takes place upon the interactions of CYP2E1-BODIPY with the membrane. To quantify the changes in the fluorescence of BODIPY we used a linear least-squares fitting of the spectra of the first and second principal components by a combination of the prototypical spectra of emission of CYP2E1-BODIPY in solution and in a membrane-bound state. These spectral standards were normalized to have the same spectral area.

The equation for the equilibrium of binary association (dimerization) used in the fitting of oligomerization isotherms (dependencies of FRET efficiency on the concentration of P450 in membranes) had the following form:

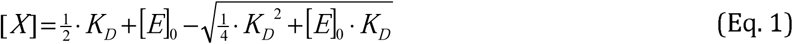

where [*E*]_0_, [*X*], and *K*_D_ are the total concentration of the associating compound (enzyme), the concentration of its dimers, and the apparent dissociation constant, respectively. In order to use this equation in fitting of the dependencies of the relative increase in fluorescence observed at enzyme concentration [*E*]_0_ (*R*_E_) equation (1) was complemented with the parameter *R*_max_. This parameter corresponds to the value of *R*_E_ observed upon a thorough dissociation of completely oligomerized enzyme:

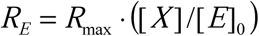

The parameter *R*_max_ is determined by the efficiency of FRET (*E*_FRET_) according to the following relationship:

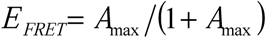

Fitting of the titration isotherms to the above equations was performed with non-linear regression using a combination of Nelder-Mead and Marquardt algorithms as implemented in our SpectraLab software [34].

## Results

### Characterization of HLM preparations used in this study

Both samples of HLM used in this study, namely HLM-1 and HLM-2, were characterized with measuring the total content of cytochromes P450 and the content of cytochrome *b*_5_ with UV-Vis spectroscopic assays. We also determined the content of CYP2E1, CYP1A2, CYP3A4 and NADPH-cytochrome P450 reductase in both samples with mass-spectrometry. The concentration of NADPH-cytochrome reductase (CPR) was also estimated on the basis of the rate of NADPH-dependent reduction of cytochrome c. Notably, the results of the two different methods of CPR quantification gave nearly identical results with both preparations of HLM.

Results of our analysis are summarized in Table 1. As seen from this table the two samples were significantly different in the composition of the P450 pool, as well as in the P450-to-phospholipids and P450-to-reductase molar ratios. HLM-2 preparation is characterized with higher P450 concentration (as judged from the phospholipids:P450 molar ratio, *R*_L/P_) and has a higher molar excess of P450 over the CPR. At the same time HLM-1 has higher content of both CYP1A2 and CYP2E1, while it is characterized with considerably lower abundance of CYP3A4 (Table 1).

**Table 1.** Characterization of microsomal preparations used in this study.

| Parameter | HLM-1 | HLM-2 |
| --- | --- | --- |
| Total P450, nmol/mg protein | 0.256±0.020 | 0.398±0.053 |
| CYP2E1, nmol/mg protein | 0.032 | 0.028 |
| CYP1A2, nmol/mg protein | 0.032 | 0.017 |
| CYP3A4, nmol/mg protein | 0.057 | 0.219 |
| CPR, nmol/mg (activity assay) | 0.058±0.010 | 0.051±0.012 |
| CPR, nmol/mg (MS assay) | 0.059 | 0.051 |
| b<sub>5</sub>, nmol/mg protein | 0.375±0.019 | 0.329±0.002 |
| Phospholipid (PL), nmol/mg protein | 724±136 | 650±136 |
| PL/P450 molar ratio | 2828 | 1633 |
| P450/CPR molar ratio | 4.4 | 7.7 |
| CYP2E1 molar fraction<sup>a</sup>, % | 12.5 | 7.1 |
| CYP1A2 molar fraction<sup>a</sup>, % | 12.7 | 4.3 |
| CYP3A4 molar fraction<sup>a</sup>, % | 22.4 | 55.1 |

\* The estimates obtained from spectroscopic and enzyme activity assays represent the averages of 2-8 individual measurements. The “±” values are the confidence intervals calculated for $p=0.05$ . The values obtained with mass-spectroscopy are the results a single measurement.
<sup>a</sup> The molar fraction was calculated from the ratio of the content of the individual P450 determined with mass spectrometry to the total P450 content determined with absorbance spectroscopy.

### Incorporation of CYP2E1 into HLM and its effect on homo-oligomerization of the protein

In our previous studies we demonstrated that the incubation of either the microsomes from insect cells (Supersomes™ or Baculosomes®) or HLM with purified CYP3A4, CYP2D6 or CYP2E1 results in efficient incorporation of the heme proteins into the membrane [9, 17, 20]. In order to further refine and better characterize this model system we developed a new technique for monitoring the process of CYP2E1 incorporation into the membrane. Our method utilizes fluorescence resonance energy transfer (homo-FRET) between fluorophores attached to neighboring CYP2E1 molecules in the oligomer. Short Stockes shift characteristic to BODIPY-based fluorescent probes results in an efficient FRET between closely located fluorophores of the same kind (homo-FRET). For our experiments we chose BODIPY-577/618 maleimide due to a minimal spectral overlap of its emission band (λ_max_=618 nm) with the absorbance bands of the P450 heme. With this florescent probe the effect of homo-FRET between two BODIPY moieties (*R*_0_=53 Å) is expected to predominate over the effect of FRET from BODIPY to the P450 heme groups (*R*_0_=40 Å).

Labeling of purified CYP2E1 with BODIPY 577/618 maleimide taken in the 1:1 – 2:1 molar ratio to the protein causes neither formation of the inactivated P420 state of the protein nor any detectable change in the P450 spin equilibrium. The FRET experiments reported here were performed with CYP2E1-BODIPY conjugates bearing two BODIPY fluorophores per one protein molecule.

The intensity of fluorescence of BODIPY 577/618 incorporated into CYP2E1 is heavily quenched in the oligomers in solution due to homo-FRET. Incorporation of the protein into the microsomal membrane is expected to cause an ample increase in fluorescence due to dissociation of the protein homo-oligomers and formation of the complexes with other P450 species, where homo-FRET is not observed. Studying the dependence of the amplitude of this effect on the amount of the incorporated protein allows to determine the apparent dissociation constant of the homo-oligomers. Furthermore, comparison of the results obtained in HLM with those derived from the studies with model microsomes (Supersomes™) containing no P450 proteins or bearing a single recombinant P450 enzyme may be used to probe the interactions of CYP2E1 with individual P450 species.

A series of spectra of fluorescence taken in the process of interactions of CYP2E1-BODIPY with HLM shown in Fig 1a demonstrate a multifold increase in the intensity of emission in full compliance with the expectations. Inset in Fig. 1a shows a series of kinetic curves of this increase recorded at various molar ratios of phospholipids of added HLM to CYP2E1-BODIPY. These time dependencies may be approximated with a three-exponential equation, where the characteristic times of the three phases amounts to 3 ± 0.5, 47 ± 20 and 600 ± 120 min. The maximal amplitude of the increase in fluorescence estimated from this approximation increases together with a decrease in the concentration of CYP2E1 in the membrane (or with an increase in the lipid-to-CYP2E1 ratio, *R*_L/P_). This observation suggests that CYP2E1 incorporation into HLM results in dissociation of the homo-oligomers present in solution and the degree of this dissociation increases with decreasing the concentration of CYP2E1 in the membrane.

**Fig.1.**
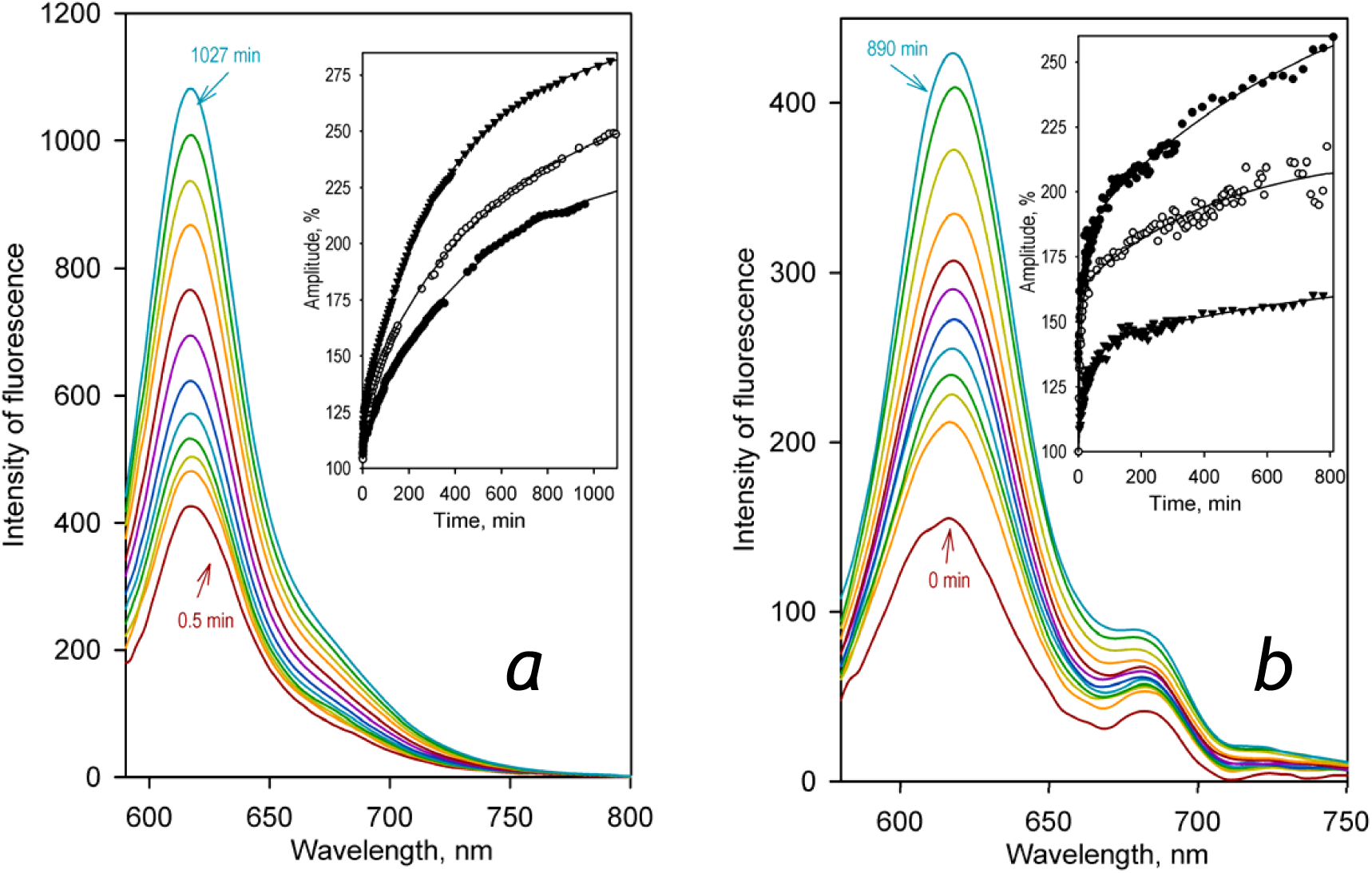
Effect of incorporation of CYP2E1-BODIPY into HLM-1 (***a***) and SS-RR (***b***) on the intensity of BODIPY fluorescence. The main panels show the series of spectra recorded in the process of incorporation and the inserts show the kinetic curves of the increase in the relative intensity of fluorescence of the probe. The spectra shown in panel ***a*** were recorded at 0.5, 2, 5, 16, 33, 67, 120, 230, 370, 530, 740 and 1027 min after addition of CYP2E1-BODIPY to HLM-1 to reach the CYP2E1-to-phospholipids molar ratio (*R*_L/P_) of 560:1 (or 0.33 µM CYP2E1-BODIPY per mg of microsomal protein). Kinetic curves shown in the inset were recorded at *R*_L/P_ ratios of 1:213 (filled circles), 1:560 (open circles) and 1:2240 (filled triangles). The spectra shown in panel ***b*** were recorded at 0.1, 0.6, 3,6, 11, 24, 49, 95, 190, 390,780 and 890 min after addition of CYP2E1-BODIPY to SS-RR to reach the *R*_L/P_ ratio of 3600:1. Kinetic curves shown in the inset were recorded at *R*_L/P_ ratios of 1:600 (filled triangles), 1:560 (open circles) and 1:2240 (filled circles).

Similar effect was also observed during the incorporation of CYP2E1-BODIPY into the membrane of control Supersomes™ (SS-RR) containing recombinant CPR, but having no cytochromes P450 (Fig 1B). In this case the dependence of the amplitude on R_L/P_. was considerably more rapid than that observed with HLM. This difference indicates that the interactions with cytochromes P450 present in HLM membrane keep CYP2E1-BODIPY away from its homo-oligomerization.

The dependencies of the amplitude of fluorescence increase on the surface density of CYP2E1-BODIPY in the membrane are shown in Fig. 2. Similar to the approach used in our previous studies [9, 17, 20] we determined the value of the apparent dissociation constant of the homo-oligomers (*K*_D_) from the fitting of these dependencies to the equation for the equilibrium of binary association (dimerization) (Eq. 1), as described in Materials: and Methods.

**Fig. 2.**
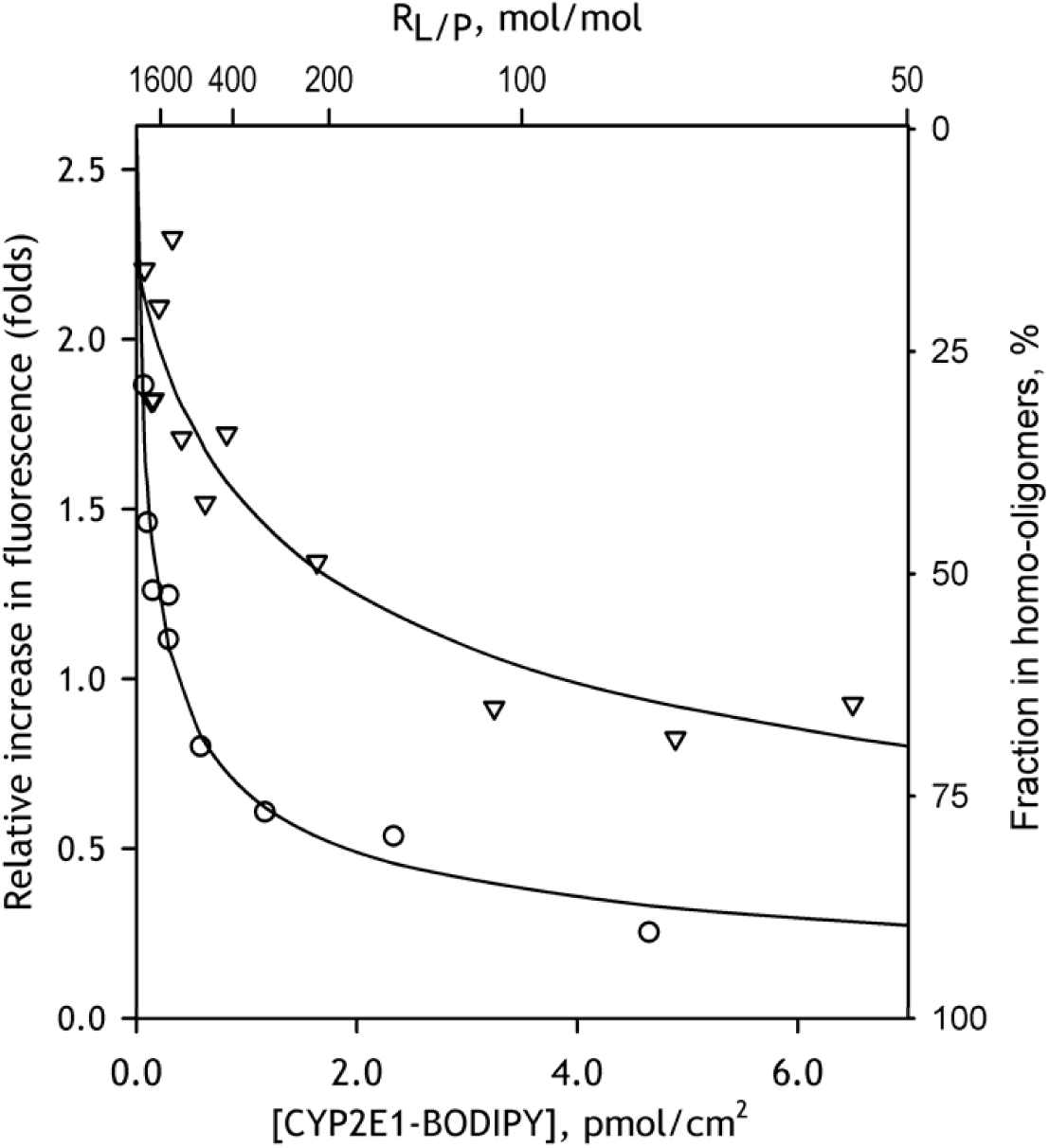
The dependencies of the relative increase in fluorescence intensity caused by incorporation of CYP2E1-BODIPY into the membranes of HLM-1 (triangles) and SS-RR (circles) on the concentration (surface density) of CYP2E1-BODIPY in the membrane. Solid lines show the approximations of the data sets with Eq. 1. The scale shown on the right of the plot indicates the apparent fraction of CYP2E1 present in homo-oligomeric state estimated from the results of this fitting.

The parameters of CYP2E1 interactions with HLM and SS-RR determined from the dependencies shown in Fig. 2 may be found in Table 2. This table also contains the parameters obtained in the studies of interactions of CYP2E1-BODIPY with CYP1A2 and CYP2C19 (see infra). As seen from this table the value of *K*_D_ for CYP2E1 homo-oligomers is much lower than that observed with the control microsomes. Therefore, the effect of CYP2E1 incorporation on its homo-oligomerization is much better pronounced in HLM as compared to the microsomes containing no P450 proteins. This observation evidences the formation of mixed complexes of CYP2E1 with other P450 species that takes place in the expense of dissociation of CYP2E1 homo-oligomers.

**Table 2.** Effect of incorporation of CYP2E1-BODIPY on the degree of its homo-olegomerization explored with homo-FRET*.

| Microsomal preparation | [CYP2E1]<sub>50%</sub><sup>a</sup>,<br>pmol/cm<sup>2</sup> | (Lipid/CYP2E1)<sub>50%</sub><sup>b</sup> | FRET<br>Efficiency, % |
| --- | --- | --- | --- |
| SS(CPR) | 0.081 ±0.020 | 4323 ±1113 | 72.9 |
| SS(b5 and CPR) | 0.143 ±0.038 | 2444 ±699 | 71.8 |
| HLM | 0.914 ±0.245 | 383 ±111 | 79.0 |
| SS(1A2) | 0.890 ±0.040 | 393 ±18 | 78.6 |
| SS(3A4) | 0.900 ±0.204 | 388 ±93 | 77.9 |
| SS(2C19) | 0.059 ±0.021 | 5891 ±2441 | 80.0 |

\* The values given in the Table were obtained by fitting of the titration curves with the binary association equation (Eq. 1). The “±” values show the confidence interval calculated for $p = 0.05$ .
<sup>a</sup> Surface density of P450 in microsomal membranes at which the amplitude of the titration curves reaches 50% of the maximal level. This value is equal to the value of $K_D$ determine from the fitting of the titration curves with Eq. 1.
<sup>b</sup> Lipid-to-P450 ratio at which the amplitude of the titration curves reaches 50% of the maximal level.

### Effect of enriching HLM with CYP2E1 on the metabolism of CYP2E1-specific substrates

In order to probe if the incorporation of CYP2E1 into HLM makes the exogenous enzyme a fully functional member of the P450 ensemble we probed the effect of enriching HLM with CYP2E1 on the metabolism of CYP2E1-specific substrates, *p*-Nitrophenol (pNP) and Chlorzoxazone (CLZ). In our studies with pNP we compared two different preparations of HLM, HLM-1 and HLM-2. The two preparations exhibited closely similar parameters of pNP metabolism and their dependencies on CYP2E1 concentration. With both preparations the dependencies of the reaction rate on pNP concentration revealed a noticeable homotropic cooperativity and obeyed the Hill equation with the Hill coefficient equal to 1.5±0.5 and 1.3±0.2 in HLM-1 and HLM-2 respectively.

As shown in Fig. 2, where we compare the profiles of oxidation of pNP and CLZ obtained with HLM with different CYP2E1 concentrations, enrichment of HLM with CYP2E1 results in a multifold increase in the rates of metabolism of both substrates. As illustrated in Fig. 3, the apparent *k*_cat_ derived from these titrations remains proportional to the amount of CYP2E1 in the membrane up to the content of 0.3-0.4 nmol/mg protein (or about 50% CYP2E1 in the P450 pool) and stabilizes (pNP) or decreases (CLZ) after that point. Other kinetic parameters (*K*_M_ (CLZ) or *S*_50_ and the Hill coefficient (pNP)) do not show any statistically significant changes (Table 3). These results demonstrates that the incorporated CYP2E1 becomes a fully-functional member of the P450 ensemble and do not exhibit any detectable functional differences with the endogenous CYP2E1 in HLM.

**Table 3.** Effect of incorporation of CYP2E1 into HLM on the kinetic parameters of oxidation of *p*-Nitrophenol and Chlorzoxazone*.

| Molar ratio of added CYP2E1 to P450 present in HLM | p-Nitrophenol<sup>a</sup> |  |  | Chlorzoxazone<sup>b</sup> |  |
| --- | --- | --- | --- | --- | --- |
| | $S_{50}$, $\mu\text{M}$ | N | $V_{\max}$, $\text{min}^{-1}$ | $K_M$, $\mu\text{M}$ | $V_{\max}$, $\text{min}^{-1}$ |
| 0 | 42.3 ± 7.8 | 1.5 ± 0.5 | 5.0 ± 1.9 | 54.4 ± 16.1 | 11.8 ± 2.7 |
| 0.25 | 25.8 ± 2.7 | 1.4 ± 0.6 | 9.9 ± 3.2 | 78.5 ± 29.0 | 41.9 ± 11.3 |
| 0.5 | 33.5 ± 8.7 | 1.4 ± 0.1 | 22.8 ± 11.8 | 54.2 ± 4.4 | 41.6 ± 4.3 |
| 1 | 33.0 ± 9.7 | 1.9 ± 0.8 | 17.5 ± 8.6 | 65.2 ± 20.2 | 77.9 ± 8.2 |
| 1.75 – 2.0<sup>c</sup> | 35.5 ± 12.6 | 2.0 ± 0.7 | 15.2 ± 4.6 | 86.7 ± 18.2 | 54.1 ± 11.0 |

\* The estimates given in the table represent the averages of 2-4 individual measurements. The “ ± ” values are the confidence intervals calculated for $p=0.05$ .
<sup>a</sup> The parameters of pNP metabolism given in this table were obtained with the HLM-1 preparation.
<sup>b</sup> The parameters of CLZ metabolism were obtained with the HLM-2 preparation.
<sup>c</sup> The values given in this row for the parameters of pNP hydroxylation were measured at a two-fold molar excess of CYP2E1 over the internal P450 content, while in the case of chlorzoxazone this ratio was equal to 1.75.

**Fig. 3.**
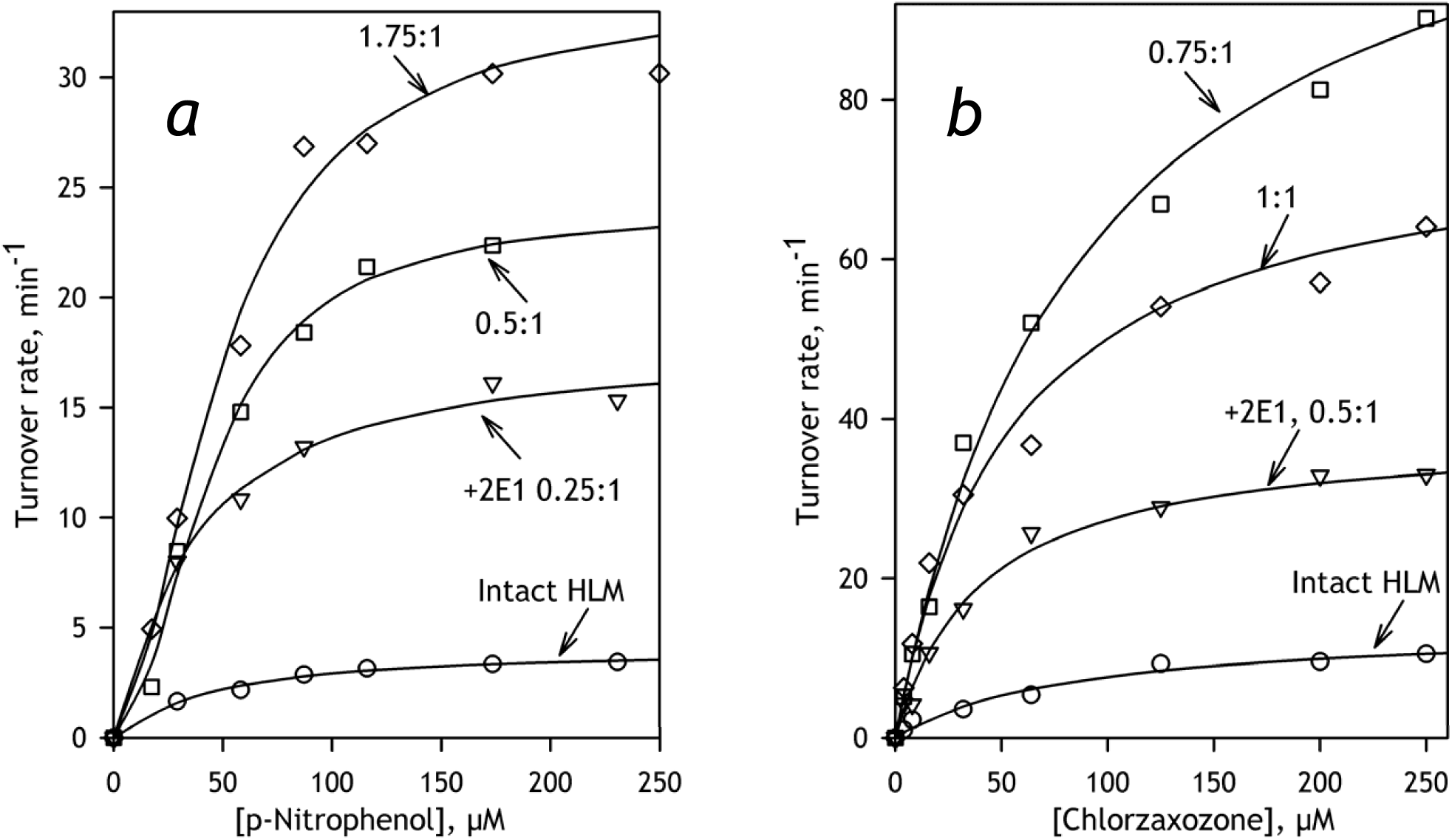
The effect of incorporation of CYP2E1 into HLM-2 on CYP2E1-dependent hydroxylation of pNP (***a***) and CLZ (***b***). The lines represent the approximations of the data sets with the Hill (***a***) or Michaelis-Menten (***b***) equations. The ratios shown next to each curve indicate the molar ratio of added CYP2E1 to the internal cytochrome P450 content in HLM-2 (0.4 nmol/mg protein).

### Probing the ability of recombinant human P450 species to metabolize CEC and ERR

In our further studies of the effect of added CYP2E1 on the metabolism of substrates of other P450 species we used fluorogenic substrates 7-etoxy-4-cyanocoumarin (CEC) and 7-ethoxyresorufin (ERR), which are primarily metabolized in HLM by CYP1A2 and CYP2C19 [35-36]. In order to better characterize the ability of human cytochrome P450 species to metabolize these substrates we investigated their metabolism by recombinant human CYP1A2, CYP2B6, CYP2C8, CYP2C9, CYP2C19, CYP2D6, CYP2E1, and CYP3A4 using commercial preparations of Supersomes™ containing the respective enzymes. Results of these assays are shown in Table 4. The probed P450 species not found in this table did not exhibit any detectable activity with either substrate.

**Table 4.** Parameters of metabolism of 7-CEC and 7-ERR by recombinant human cytochromes P450*.

| P450 species | CEC |  | ERR |  |
| --- | --- | --- | --- | --- |
| | $K_M$, $\mu\text{M}$ | $V_{\max}$, $\text{s}^{-1}$ | $K_M$, $\mu\text{M}$ | $V_{\max}$, $\text{s}^{-1}$ |
| CYP1A2 | 2.3 ±0.10 | 7.9 ±2.2 | 0.40 ±0.14 | 1.1 ±0.3 |
| CYP2B6 | 8.2 ±2.4 | 1.0 ±0.2 | No activity detected | |
| CYP2C8 | No activity detected | | 0.26 ±0.14 | 0.02 ±0.01 |
| CYP2C19 | 24 ±5 | 2.3 ±0.5 | 0.33 ±0.13 | 0.06 ±0.02 |
| CYP2E1 | No activity detected | | 33.0 ±9.2 | 0.05 ±0.02 |

\* The estimates given in the table represent the averages of 2-4 individual measurements. The “ ± ” values are the confidence intervals calculated for $p=0.05$ .

In a good agreement with the previous reports the highest activity in dealkylation of ERR was exhibited by CYP1A2. Some activity with this substrate was also detected with CYP2C8 and CYP2C19. However, although affinity of these CYP2C enzymes to ERR was quite close to that specific to CYP1A2, the rates of reaction observed with them were considerably lower. Although CYP2E1 also reveals some activity in ERR dealkylation, the observed values of *V*_max_ and *K*_M_ (Table 4) leave out a possibility of any sizeable participation of CYP2E1 in ERR metabolism by HLM, at least at the substrate concentrations below 5 µM used in our studies with CYP2E1-enriched HLM.

Although the highest rate of metabolism of CEC was also exhibited by CYP1A2, the activity rendered by CYP2C19 is high enough to consider this enzyme as an important contributor to CEC metabolism, especially taking into account its relatively high abundance in HLM [4]. Another P450 enzyme capable of metabolizing CEC is CYP2B6. However, this enzyme is unlikely to make any important contribution to CEC metabolism in HLM due to low rate of turnover and low concentration of CYP2B6.

Therefore, probing the impact of CYP2E1 incorporation on the parameters of metabolism of CEC and ERR by HLM is expected to provide information on the activities of CYP1A2 and CYP2C19 and discriminate between the CYP2E1 effects on the two enzymes.

### Impact of CYP2E1 incorporation on the metabolism of CEC and ERR by HLM

The dependencies of the rate of CEC dealkylation on substrate concentration obtained with intact and CYP2E1-enriched HLM samples HLM-1 and HLM-2 are shown in Fig. 5. The parameters obtained from the approximation of these dependencies with the Michaelis-Menten equation are summarized in Table 5. The rate of metabolism of CEC by HLM-1 was noticeably higher than that observed with HLM-2, consistent with higher concentration of CYP1A2 in HLM-1 as compared to HLM-2 (Table 1). As suggested by the titration traces shown in Fig. 5, CYP2E1 enrichment may result in some increase in the rate of CEC metabolism in both preparations of HLM. However, this increase cannot be considered statistically significant (Table 5) due to a large bias of the results of individual measurements. More important is a statistically significant decrease in the value of *K*_M_ observed with both HLM preparations (Table 5). While in the intact supersomes this value is close to that characteristic to CYP2C19-dependent metabolism of CEC (Table 4), incorporation of CYP2E1 results in its decrease and makes it closer to that is expected for CYP1A2-catalyzed reaction (Table 5).

**Table 5.** Parameters of CEC dealkylation in intact and CYP2E1-enriched HLM obtained from the fitting of titration curves with the Michaelis-Menten equation.

| HLM sample | $K_M$, $\mu\text{M}$ | $V_{\max}$, $\text{min}^{-1}$ |
| --- | --- | --- |
| HLM-1 | 15.6 ± 3.0 | 3.4 ± 0.3 |
| HLM-1 + CYP2E1<sup>a</sup> | 9.4 ± 1.2 (0.004) | 3.8 ± 0.3 (0.159) |
| HLM-2 | 32.2 ± 14.7 | 1.9 ± 0.5 |
| HLM-2 + CYP2E1<sup>a</sup> | 6.1 ± 1.9 (0.009) | 2.3 ± 0.8 (0.450) |

\* The estimates given in the table represent the averages of 2-4 individual measurements. The “ ± ” values are the confidence intervals calculated for $p=0.05$ . The values in parentheses represent the $p$ -values of Student's t-test for the hypothesis of equality of the respective values with the value presented in the previous line (i.e., at no CYP2E1 incorporated).
<sup>a</sup> The results obtained with the samples containing CYP2E1 incorporated at 1:1 molar ratio to the microsomal P450 content (0.26 and 0.4 nmol/mg protein for HLM-1 and HLM-2, respectively).

**Fig. 4.**
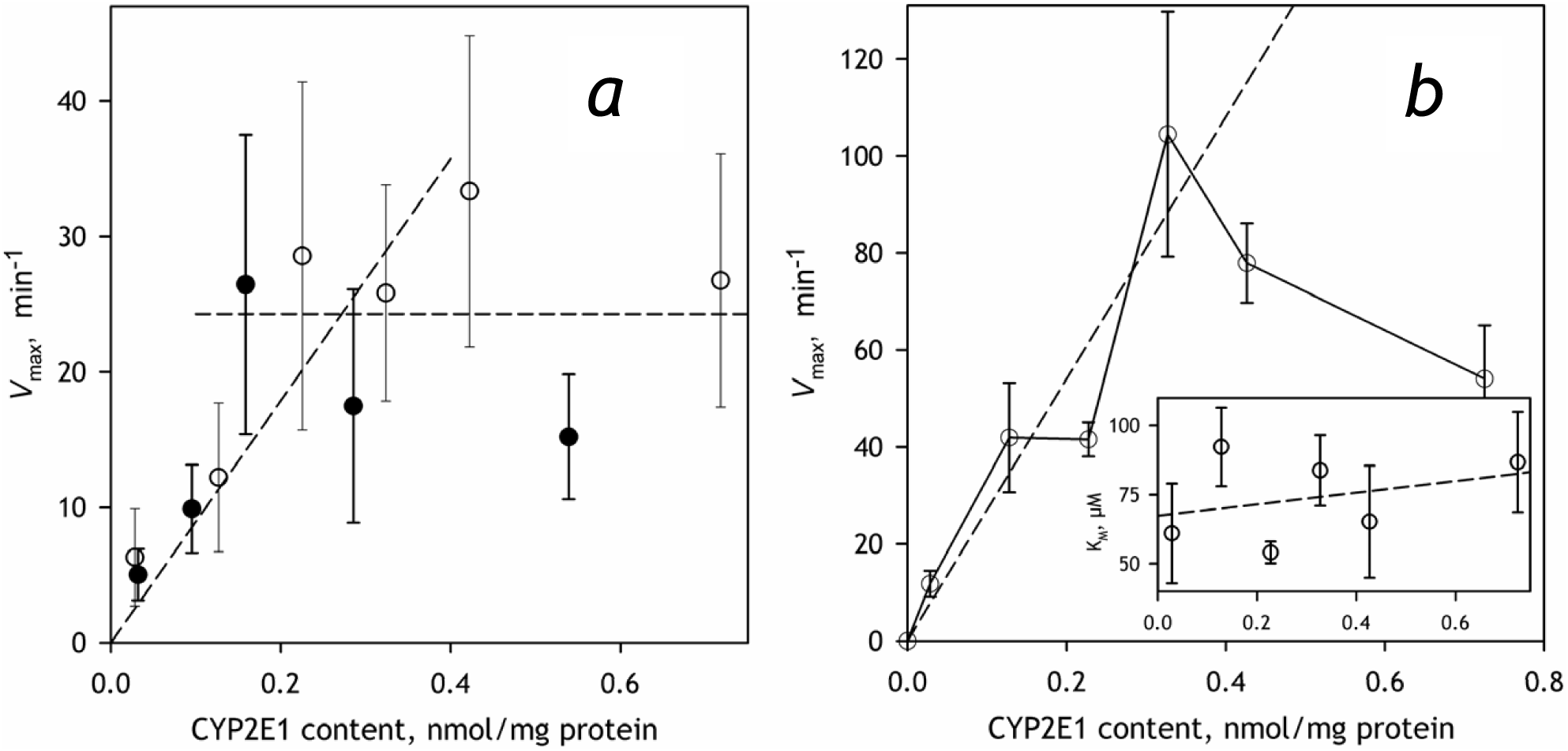
The effect of incorporation of CYP2E1 into HLM on the metabolism of CYP2E1-specific substrates pNP (***a***) and CLZ (***b***). The data shown in open and filled circles in panel ***a*** were obtained with HLM-1 and HLM-2 preparations respectively. The results presented in the panel ***b*** were obtained with HLM-2. Error bars represent the confidence intervals calculated for *p*=0.05 on the basis of 2-5 measurements.

**Fig. 5.**
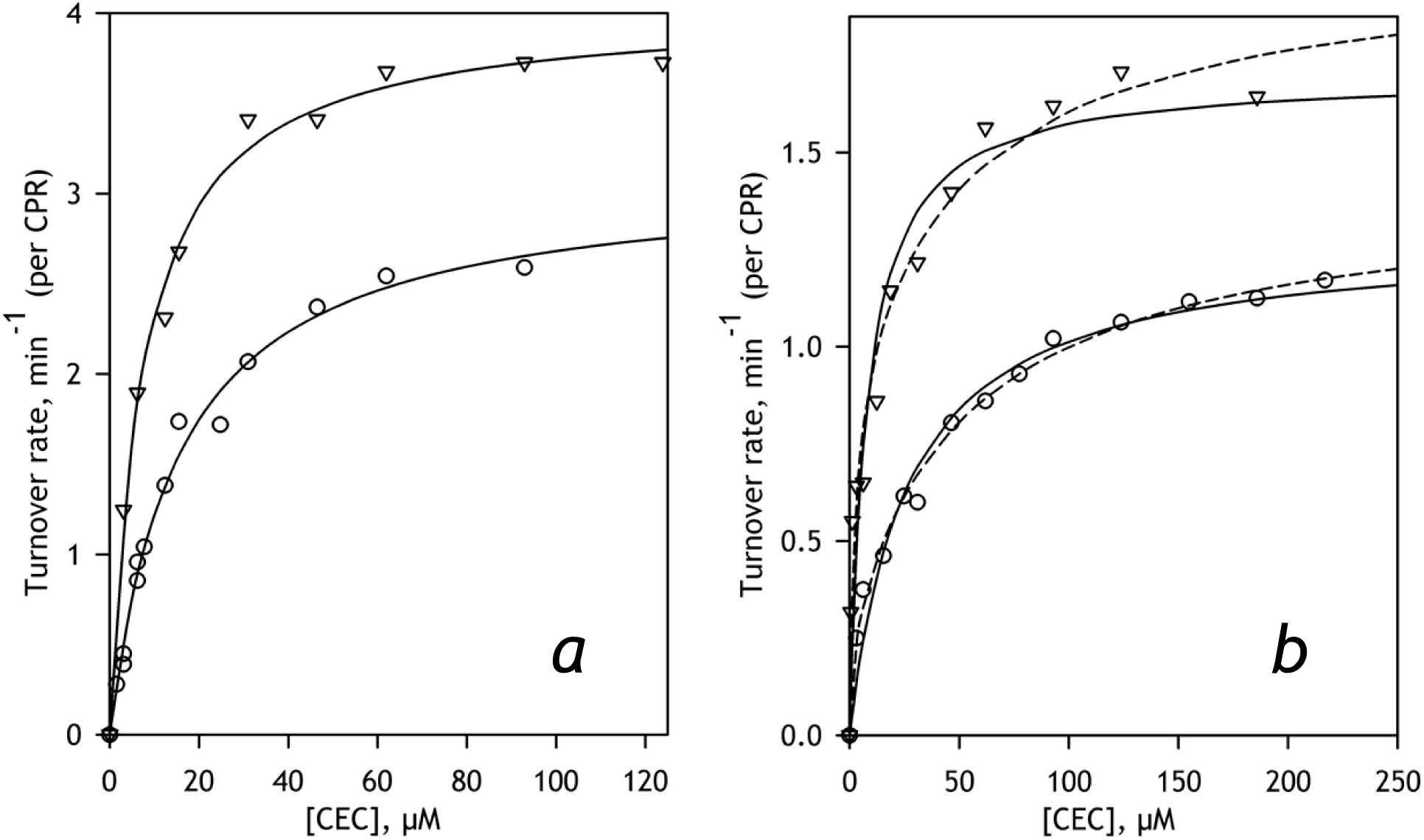
The effect of incorporation of CYP2E1 into HLM on the dependencies of the rate of CEC metabolism on substrate concentration. The results obtained with HLM-1 and HLM-2 are shown in the panels ***a*** and ***b*** respectively. The data obtained with intact HLM are shown in circles while the triangles designate the results obtained with the samples containing CYP2E1 incorporated at 1:1 ratio to the microsomal P450 content (0.26 and 0.4 nmol/mg protein for HLM-1 and HLM-2, respectively). The solid limes represent the results of fitting of the datasets to the Michaelis-Menten equation. The dashed lines in the panel ***b*** shows the approximations with the equation of the sum of two Michaelis-Menten functions.

The change in the main route of CEC metabolism suggested by this initial analysis may be better revealed through fitting the titration curves to the equation of the sum of two Michaelis-Menten functions. In the case of HLM-1, however, this fitting was incapable to resolve two components with different *K*_M_ values and inefficient in improving the quality of approximation. However, in the case of HLM-2 the fitting of the datasets with a single Michaelis-Menten function results in systematic deviations suggesting a noticeable biphasisity of the titration curves (Fig. 5b). The use of the equation of the sum of two Michaelis-Menten functions for fitting the datasets obtained with HLM-2 (either with or without incorporated CYP2E1) suggests the values of Km for the two phases to be in the range of 2-4 and 30-70 µM.

Global fitting of the entire dataset obtained with a series of HLM-2 preparations containing various amounts of incorporated CYP2E1 gives the best results with the estimates of *K*_M_ of 2.5±0.9 and 58±13 µM for the high- and low-affinity components respectively. These estimates are in a reasonable agreement with the values of 2.3±0.1 and 24±5 µM 2.3 obtained with CYP1A2- and CYP219-containing Supersomes respectively (Table 4). As illustrated in Fig. 6a, the fraction of the high-affinity phase derived from the global fitting increases proportionally to the concentration of CYP2E1 in HLM up to the content of 0.2-0.3 nmol/mg protein and tends to stabilize after that point. This change is likely to be also associated with some increase in the rate of CEC metabolism, although the latter effect cannot be considered statistically significant (Fig. 5b).

**Fig. 6.**
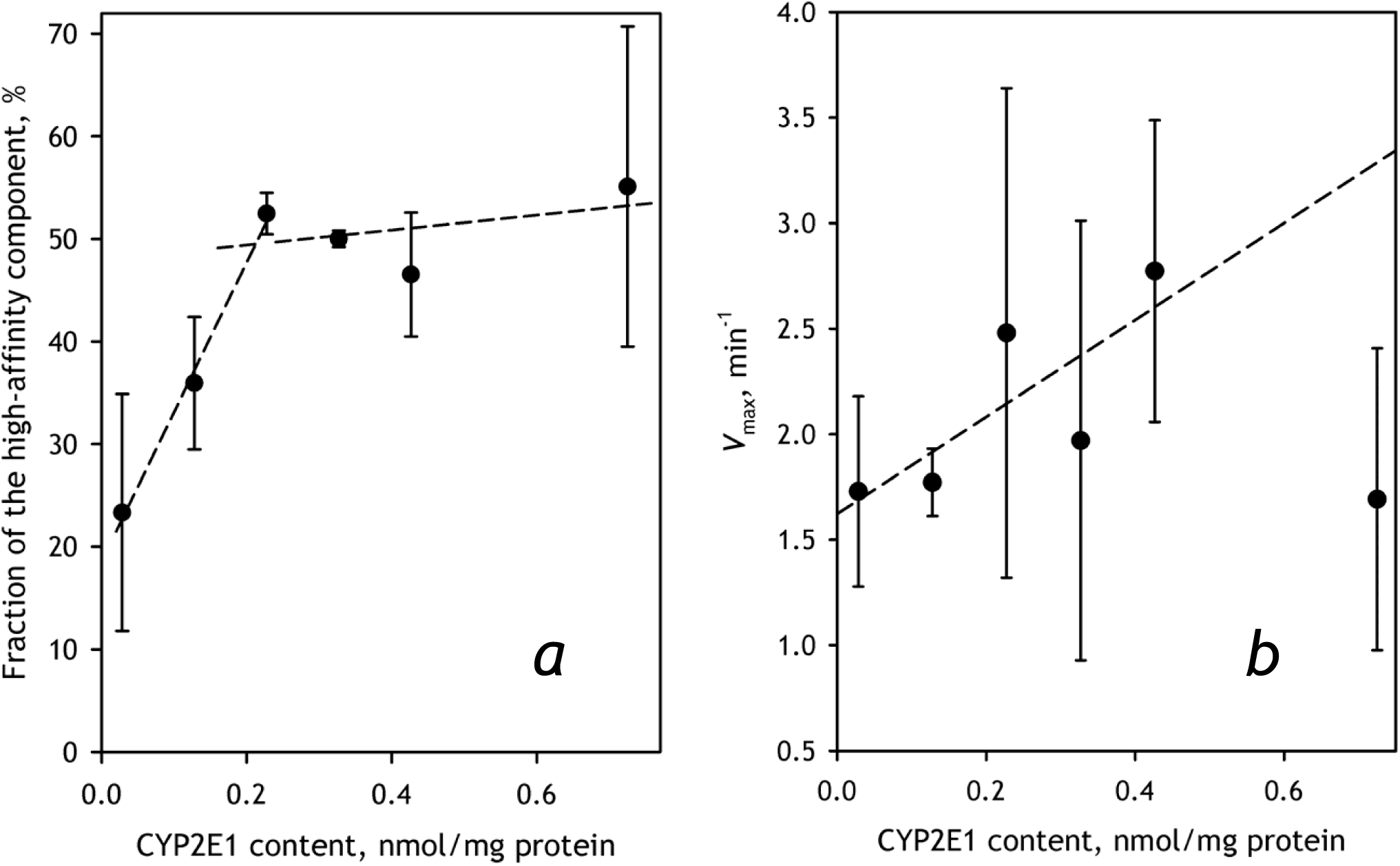
The effect of incorporation of CYP2E1 into HLM-2 on the parameters of metabolism of CEC. Panel ***a*** shows the dependence of the fraction of the high-affinity component on CYP2E1 content. The dependence of the total maximal rate of metabolism is sown in panel ***b***. Error bars represent the confidence intervals calculated for *p*=0.05 on the basis of 2-4 measurements.

The effect of CYP2E1 enrichment on the contribution of the high affinity component in the biphasic profile of CEC dealkylation suggests an important augmentation of the involvement of CYP1A2. In order to probe this possible activation further we investigated the effect of incorporation of CYP2E1 on the metabolism of ERR, the substrate almost exclusively metabolized by CYP1A2. As shown in Fig. 6 increase in the concentration of CYP2E1 causes a remarkable increase in the rate of ERR dealkylation up to the CYP2E1 content of 0.4 nmol/mg protein (or about 50% CYP2E1 in the P450 pool). The fact that this multifold increase in *V*_max_ is not associated with any substantial changes in the value of *K*_M_ (Fig. 6b, inset) rules out a possibility of direct involvement of CYP2E1-dependent metabolism of CEC and substantiates our conclusion that incorporation of CYP2E1 results in an ample activation of CYP1A2 present in HLM.

### Probing the association of CYP2E1 with CYP1A2 and CYP2C19 in Supersomes

In order to probe if the observed activation of CYP1A2 may be attributed to a direct consequence of the formation of mixed oligomers of CYP1A2 and CYP2E1 we probed the association of these two proteins in the microsomal membrane using CYP1A2-containing Supersomes™ and our FRET-based technique that utilizes CYP2E1 labeled with BODIPY 577/618. The same approach was also used to probe possible interactions of CYP2E1 with CYP2C19, another enzyme metabolizing CEC. In order to have an appropriate control dataset we also monitored the interactions of CYP2E1 with Supersomes™ containing human CPR and cytochrome b_5_ but bearing no P450 enzymes.

Similar to what was observed upon incorporation of CYP2E1-BODIPY into HLM (Fig. 1), addition of CYP2E1-BODIPY to the control Supersomes™ as well as to those containing CYP1A2 or CYP2C19 resulted in a stepwise increase in the intensity of fluorescence that obeyed three-exponential kinetics and took 18 – 20 hours for completion. The dependencies of the maximal amplitude of this increase on CYP2E1 concentration in the membrane (Fig. 7) demonstrate a significant difference between the CYP1A2-containing microsomes with the microsomes bearing no P450 protein or having CYP2C19 in their membranes. In the latter two cases a substantial degree of dissociation of the CYP2E1-BODIPY homo-oligomers was observed only at low surface densities of the incorporated enzyme, at *R*_L/P_ ratio in the range of several thousands. In contrast, in the case of CYP1A2 containing microsomes 50% dissociation of homo-oligomers was observed at *R*_L/P_ ratio around 400:1, similar to what was seen in HLM (Table 2). These results suggest that CYP2E1 efficiently forms mixed oligomers with CYP1A2 but lacks any ability to interact with CYP2C19, which stands in one-to-one correspondence with the observed effect of CYP2E1 on CYP1A2-specific activities in HLM.

**Fig. 7.**
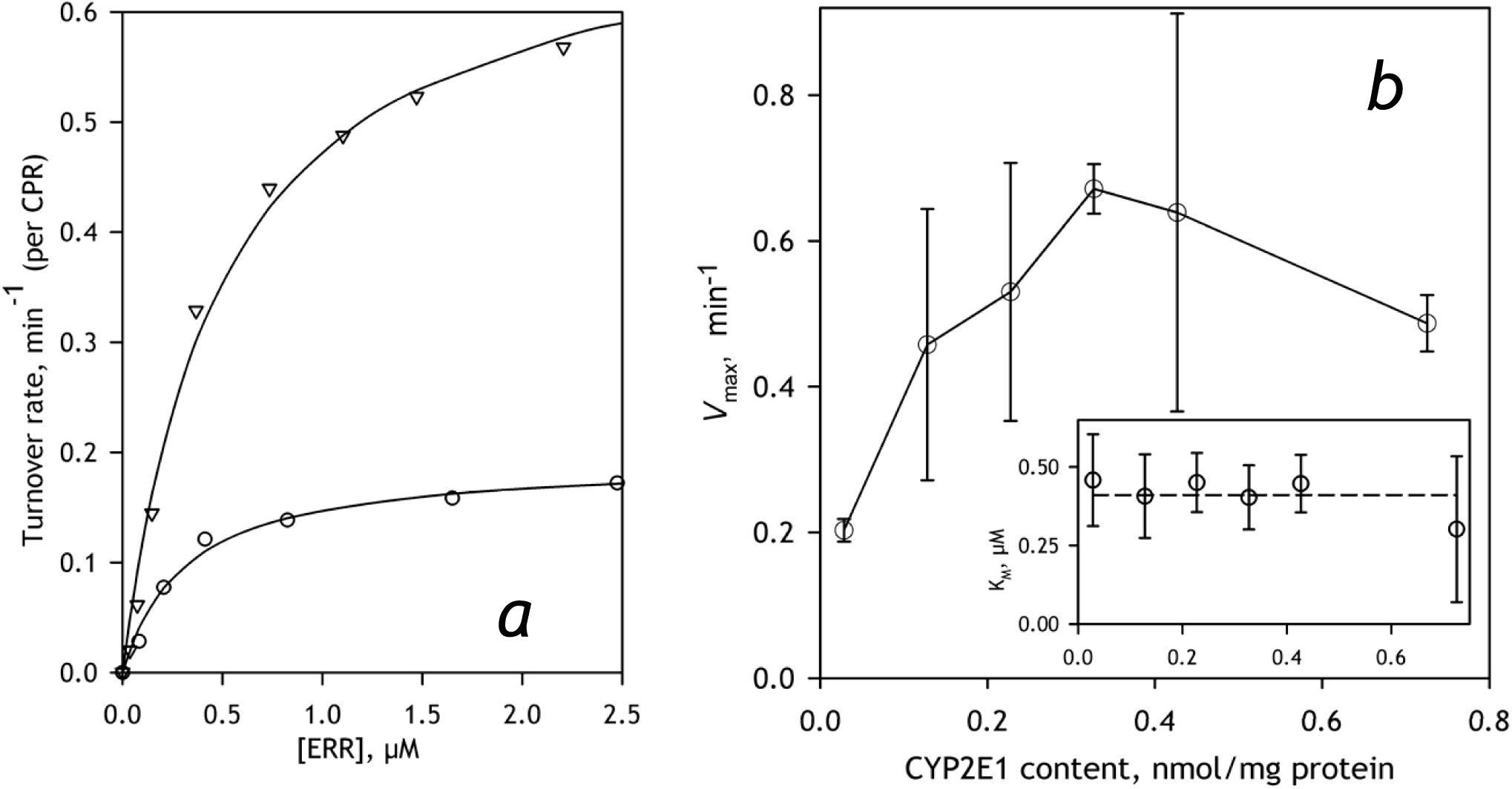
The effect of incorporation of CYP2E1 into HLM-2 on the parameters of metabolism of ERR. Panel ***a*** exemplified the dependence of the rate of turnover on substrate concentration obtained with intact HLM (circles) and the sample containing CYP2E1 incorporated in the amount of 0.3 nmol per mg protein that corresponds to 0.75:1 ratio to the microsomal P450 content (triangles). Panel ***b*** shows the dependencies of *V*_max_ (main panel) and *K*_M_ on CYP2E1 content. Error bars represent the confidence intervals calculated for *p*=0.05 on the basis of 2-5 measurements.

## Discussion

In this study we utilized the approach based on homo-FRET (fluorescent resonance energy transfer where donor and acceptor are represented by the fluorophores with one and the same structure) to monitor the incorporation of CYP2E1 into HLM and probe its interactions with other cytochrome P450 species in the microsomal membrane. Studying the effect of this incorporation on CYP2E1-specific activities of HLM we demonstrate that the incorporated enzyme becomes a fully-functional member of the P450 ensemble and do not exhibit any detectable functional differences with the natively-present enzyme. Exploring the functional effect of increase in CYP2E1 concentrations we evidenced an important activation of CYP1A2 that results in rerouting of metabolism of CEC, the substrate jointly metabolized by CYP1A2 and CYP2C19, toward preferential dealkylation by CYP1A2. Furthermore, probing the interactions of CYP2E1 with model microsomes containing individual P450 enzymes we found that CYP2E1 efficiently interacts with CYP1A2, but lacks any ability to form complexes with CYP2C19. This observation stands in one-to-one correspondence with the observed effect of CYP2E1 on CYP1A2-specific activities in HLM.

What though may be the mechanism of activation of one P450 species by its interactions with another, dissimilar P450 enzyme observed in this study and also evidenced in earlier reports from our group and by others [9, 16-17, 37-38]? Our understanding of these instances is based on a concept of “positional heterogeneity”, which we introduced to explain a persistent distribution of P450 pool between the fractions with different functional properties [5, 8, 17]. This concept is based on an assumption that the P450 subunits forming homo- and heterooligomers are not identical in their conformation and orientation, but are characterized by different abilities to be reduced, bind substrates and interact with redox partners. In effect, oligomerizartion of cytochromes P450 results in abstracting an important part of the P450 pool from the catalytic activity. A discussion of possible structural basis for positional heterogeneity in P450 oligomers may be found in a recent comprehensive review by Reed and Backes [39].

From the standpoint of the above concept, the most obvious explanation for non-additive effects of changing the content of cytochrome P450 ensemble might be a difference in the propensities of different P450 species for occupying the positions of two different types in the oligomer. In particular, it might be hypothesized that, when CYP1A2 interacts with CYP2E1, the former preferentially occupies the “active” positions, whereas CYP2E1 fills the “hindered” ones. This re-distribution of P450 species between the positions of two different types result in increase in the active fraction of CYP1A2.

This hypothesis is illustrated in Fig. 9, which schematically represents the sequence of events taking place upon incorporation of CYP2E1 (orange subunits) into the microsomal membrane. In this figure we arbitrary depicted P450 oligomer in solution as a hexamer composed of two types of subunits – “active” (triangle-shaped) and “hindered” (diamond-shaped). The same types of subunits also present in the P450 oligomers in the membrane, which are shown in Fig. 9 as trimers. Prior to incorporation of CYP2E1 the P450 species present in HLM form heterooligomers, where certain P450 species including CYP1A2, which is shown as green subunits, preferentially occupy the “hindered” locations, while the active positions are occupied by other P450 species (blue subunits). Upon incorporation of CYP2E1 (event 1) and after a slow dissociation and reassembly of the oligomers (event 2) the “hindered” positions become preferentially occupied by CYP2E1 due to some inherent structural features of the latter. This reorganization displaces CYP1A2 to “active” positions (triangle-shaped subunits) and thereby activates the enzyme.

**Fig. 8.**
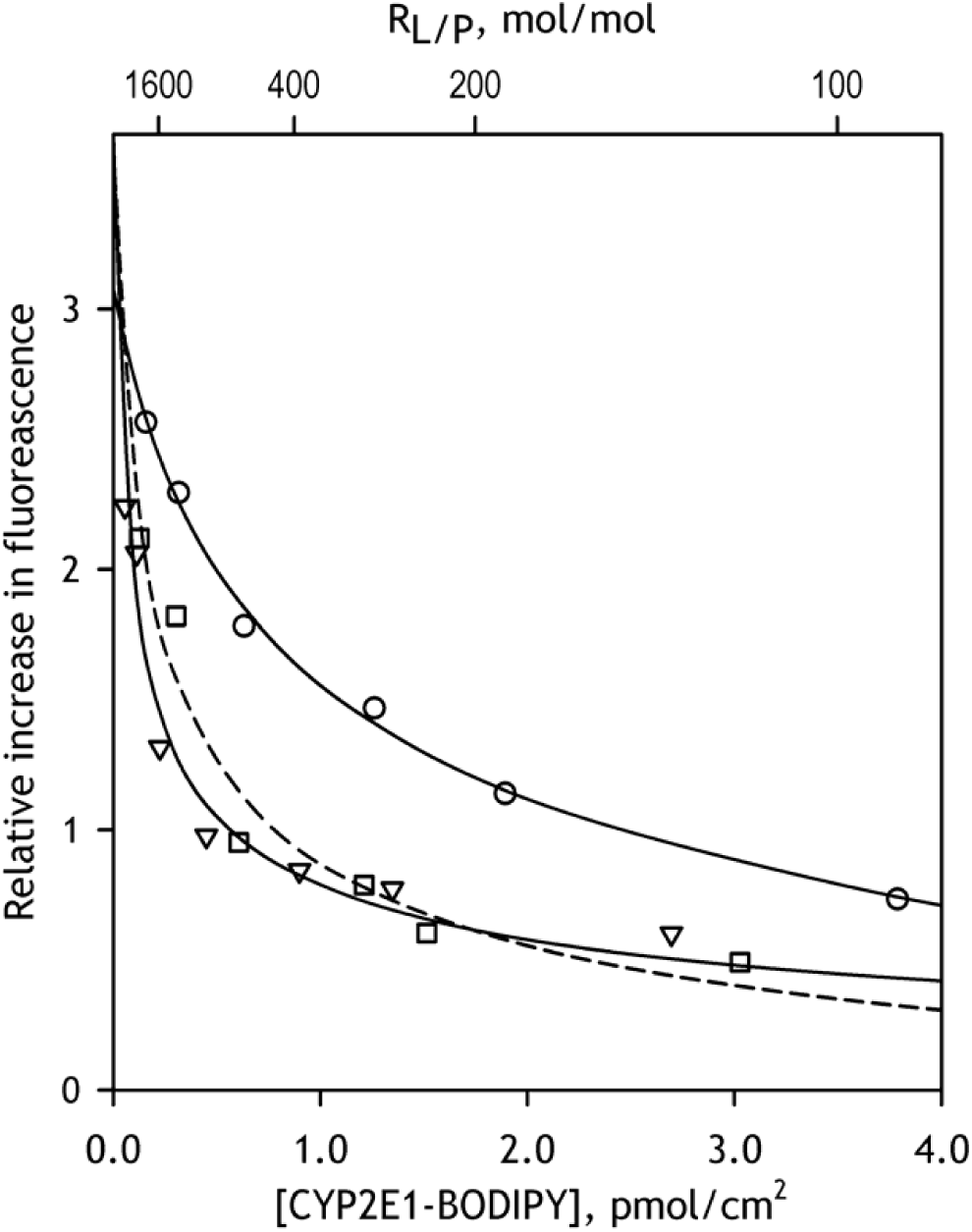
Effect of increasing concentrations of CYP2E1-BODIPY on the relative increase in fluorescence intensity caused by incorporation of CYP2E1-BODIPY into the membranes of Supersomes™ containing recombinant CYP1A2 and CPR (circles, solid line), CYP2C19, CPR and cytochrome *b*_5_ (triangles, solid line) or control Supersomes™ containing human CPR and cytochrome *b*_5_ but bearing no cytochromes P450 (squares, dashed line). The lines show the approximations of the data sets with Eq. 1.

**Fig. 9.**
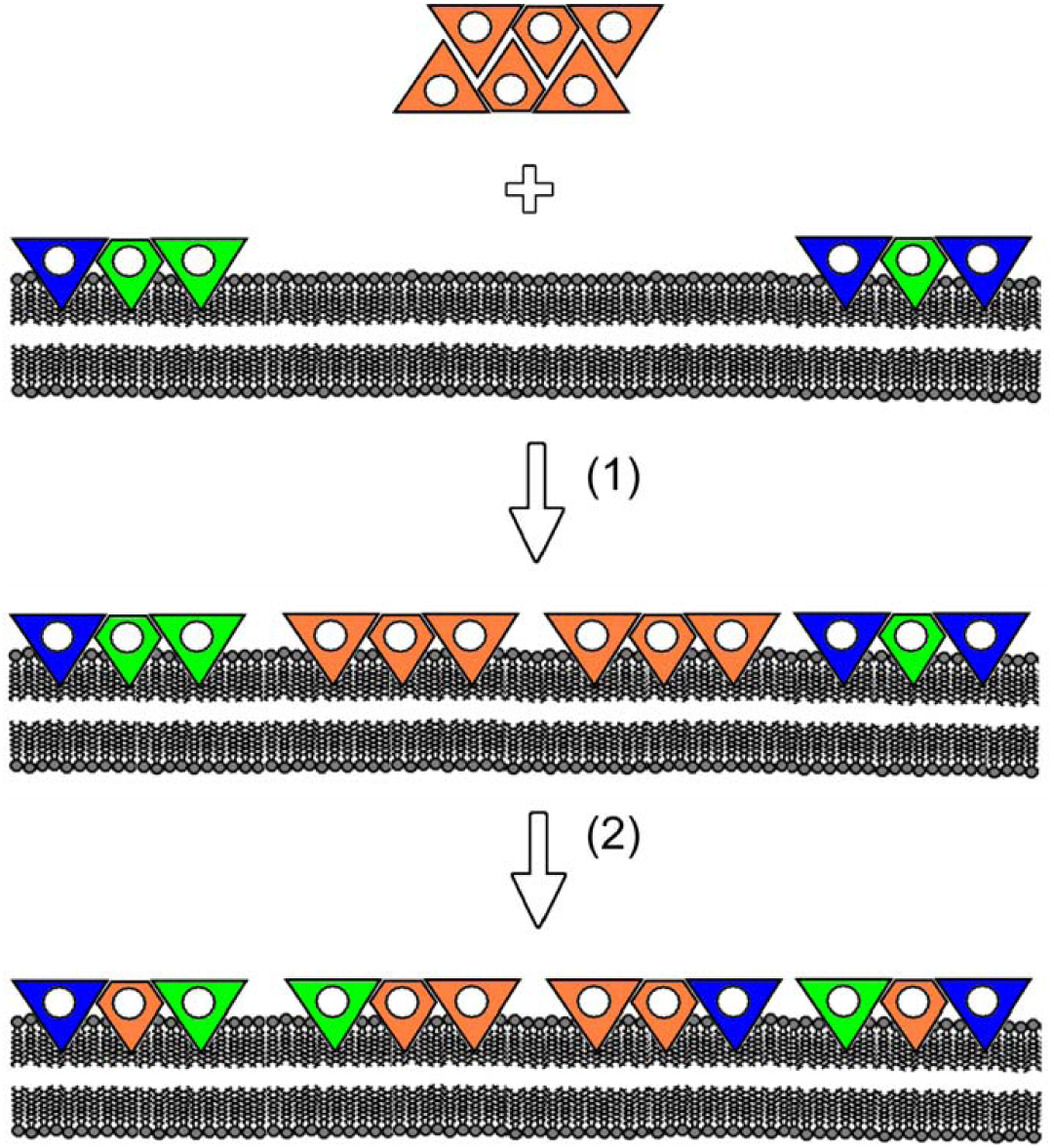
A schematic representation of the hypothesis explaining the mechanism of activation of CYP1A2 (green subunits) by CYP2E1 (orange subunits). The mechanism involves incorporation of added CYP2E1 into the membrane of HLM (event 1), where important part of CYP1A2 is rendered inactive due to a preferential occupation of the “hindered” positions (diamond-shaped subunits) in heterooligomers with other P450 species (blue subunits). After a slow dissociation and reassembly of the oligomers (event 2) the “hindered” positions become preferentially occupied by CYP2E1 due to some inherent structural features of the latter. This reorganization displaces CYP1A2 to the “active” positions (triangle-shaped subunits) and thereby activates the enzyme.

Obviously, the scheme shown in Fig. 9 is extremely simplified. It limits the number of interacting P450 species to three, does not show reversible dissociation of the oligomers and does not take into account modulation of P450 oligomerization by isoform-specific substrates discussed in our previous publication [9]. In reality the difference in the structural properties between multiple microsomal P450 species may result in very complex dependence of the degree of position-determined “hindering” of any particular P450 on the composition of the P450 pool and its exposure to P450 substrates and allosteric modulators.

The present study reveals only one aspect of this complex inter-enzyme crosstalk in the P450 ensemble – activation of CYP1A2 caused by increased content of CYP2E1 in the heterooligomers of multiple P450 species. From the practical standpoint this observation suggests that the ethanol-dependent induction CYP2E1 in human liver may have a pronounced effects on pharmacokinetics of drugs jointly metabolized by CYP1A2 and CYP2C enzymes. Further studies are planned to probe the impact of CYP2E1 on the metabolism of such major drugs as tricyclic antidepressants (amitriptylline, imipramine), and anticonvulsants (phenytoin, carbamazepine), which fall into this category.

A series of investigations with HLM preparations varying in the composition of the P450 ensemble is needed to reveal other principal elements of the entangled system of interrelations between multiple P450 species. The present study demonstrates that the approach based on incorporation of any particular purified P450 enzyme into HLM may provide a potent means for untangling this complex system and obtaining the information necessary for unambiguous prediction of the profile of drug metabolism specific to any given composition of the P450 ensemble in human liver.

## Acknowledgments

This research was supported by the National Institute On Alcohol Abuse And Alcoholism of NlH under Award Number R21AA024548. The authors are grateful to Jeffrey P. Jones (WSU) for research support, assistance in obtaining and analyzing LC-MS/MS data and continuous interest to this study. We also gratefully acknowledge the assistance of the “Human Proteome” Core Facility of the Institute of Biomedical Chemistry (IBMC, Moscow, Russia) in generating mass-spectrometry data used in characterization of HLM preparations.

## Author contributions

DRD and VGZ designed the study, DRD wrote the paper, DRD, ND, BD, NEV, and MAM conducted the experiments; DRD and VGZ analyzed the results.

