## Supplemental Table S1 for "Non-additive effects of changing the cytochrome P450 ensemble: incorporation of CYP2E1 into human liver microsomes and its impact on CYP1A2"

<u>Table S1</u>. Mass spectrometric parameters of the peptides used for quantitative analysis

| Protein Name | Peptide Modified Sequence | Precursor Mz | Precursor Charge | Collision Energy | Product Mz | Product Charge | Fragment Ion | Fragment Ion Type | Fragment Ion Ordinal |
|---|---|---|---|---|---|---|---|---|---|
| P08684,CP3A4_HUMAN | LSLGGLLQPEKPVVLK | 846.026923 | 2 | 27.6 | 1150.719421 | 1 | y10 | y | 10 |
| P08684,CP3A4_HUMAN | LSLGGLLQPEKPVVLK | 846.026923 | 2 | 27.6 | 1037.635357 | 1 | y9 | y | 9 |
| P08684,CP3A4_HUMAN | LSLGGLLQPEKPVVLK | 846.026923 | 2 | 27.6 | 909.57678 | 1 | y8 | y | 8 |
| P08684,CP3A4_HUMAN | LSLGGLLQPEKPVVLK | 846.026923 | 2 | 27.6 | 555.38646 | 1 | y5 | y | 5 |
| P08684,CP3A4_HUMAN | LSLGGLLQPEKPVVLK | 850.034022 | 2 | 27.6 | 1158.73362 | 1 | y10 | y | 10 |
| P08684,CP3A4_HUMAN | LSLGGLLQPEKPVVLK | 850.034022 | 2 | 27.6 | 1045.649556 | 1 | y9 | y | 9 |
| P08684,CP3A4_HUMAN | LSLGGLLQPEKPVVLK | 850.034022 | 2 | 27.6 | 917.590979 | 1 | y8 | y | 8 |
| P08684,CP3A4_HUMAN | LSLGGLLQPEKPVVLK | 850.034022 | 2 | 27.6 | 563.400659 | 1 | y5 | y | 5 |
| P05181,CP2E1_HUMAN | GIIFNNGPTWK | 623.83258 | 2 | 16.3 | 1076.552356 | 1 | y9 | y | 9 |
| P05181,CP2E1_HUMAN | GIIFNNGPTWK | 623.83258 | 2 | 16.3 | 963.468292 | 1 | y8 | y | 8 |
| P05181,CP2E1_HUMAN | GIIFNNGPTWK | 623.83258 | 2 | 16.3 | 816.399878 | 1 | y7 | y | 7 |
| P05181,CP2E1_HUMAN | GIIFNNGPTWK | 623.83258 | 2 | 16.3 | 702.35695 | 1 | y6 | y | 6 |
| P05181,CP2E1_HUMAN | GIIFNNGPTWK | 627.839679 | 2 | 16.3 | 1084.566555 | 1 | y9 | y | 9 |
| P05181,CP2E1_HUMAN | GIIFNNGPTWK | 627.839679 | 2 | 16.3 | 971.482491 | 1 | y8 | y | 8 |
| P05181,CP2E1_HUMAN | GIIFNNGPTWK | 627.839679 | 2 | 16.3 | 824.414077 | 1 | y7 | y | 7 |
| P05181,CP2E1_HUMAN | GIIFNNGPTWK | 627.839679 | 2 | 16.3 | 710.371149 | 1 | y6 | y | 6 |
| P16435,NCPR_HUMAN | NPFLAAVTTNR | 602.327662 | 2 | 15.2 | 732.399878 | 1 | y7 | y | 7 |
| P16435,NCPR_HUMAN | NPFLAAVTTNR | 602.327662 | 2 | 15.2 | 661.362764 | 1 | y6 | y | 6 |
| P16435,NCPR_HUMAN | NPFLAAVTTNR | 602.327662 | 2 | 15.2 | 590.32565 | 1 | y5 | y | 5 |
| P16435,NCPR_HUMAN | NPFLAAVTTNR | 602.327662 | 2 | 15.2 | 491.257236 | 1 | y4 | y | 4 |
| P16435,NCPR_HUMAN | NPFLAAVTTNR | 607.331796 | 2 | 15.2 | 742.408147 | 1 | y7 | y | 7 |
| P16435,NCPR_HUMAN | NPFLAAVTTNR | 607.331796 | 2 | 15.2 | 671.371033 | 1 | y6 | y | 6 |
| P16435,NCPR_HUMAN | NPFLAAVTTNR | 607.331796 | 2 | 15.2 | 600.333919 | 1 | y5 | y | 5 |
| P16435,NCPR_HUMAN | NPFLAAVTTNR | 607.331796 | 2 | 15.2 | 501.265505 | 1 | y4 | y | 4 |
| P05177,CP1A2_HUMAN | ASGNLIPQEK | 528.787838 | 2 | 11.4 | 898.499258 | 1 | y8 | y | 8 |
| P05177,CP1A2_HUMAN | ASGNLIPQEK | 528.787838 | 2 | 11.4 | 614.350802 | 1 | y5 | y | 5 |
| P05177,CP1A2_HUMAN | ASGNLIPQEK | 528.787838 | 2 | 11.4 | 501.266738 | 1 | y4 | y | 4 |
| P05177,CP1A2_HUMAN | ASGNLIPQEK | 528.787838 | 2 | 11.4 | 443.224873 | 1 | b5 | b | 5 |
| P05177,CP1A2_HUMAN | ASGNLIPQEK | 528.787838 | 2 | 11.4 | 556.308937 | 1 | b6 | b | 6 |
| P05177,CP1A2_HUMAN | ASGNLIPQEK | 532.794937 | 2 | 11.4 | 906.513457 | 1 | y8 | y | 8 |
| P05177,CP1A2_HUMAN | ASGNLIPQEK | 532.794937 | 2 | 11.4 | 622.365001 | 1 | y5 | y | 5 |
| P05177,CP1A2_HUMAN | ASGNLIPQEK | 532.794937 | 2 | 11.4 | 509.280937 | 1 | y4 | y | 4 |
| P05177,CP1A2_HUMAN | ASGNLIPQEK | 532.794937 | 2 | 11.4 | 443.224873 | 1 | b5 | b | 5 |
| P05177,CP1A2_HUMAN | ASGNLIPQEK | 532.794937 | 2 | 11.4 | 556.308937 | 1 | b6 | b | 6 |
